# Unmixing the Psychedelic Connectome: Brain Network Traits of Psilocybin

**DOI:** 10.1101/2025.11.17.688834

**Authors:** Krishna Prasad Bhavaraju, Natasha L. Mason, Pablo Mallaroni, Dietmar Heinke, Stefan W. Toennes, Johannes G. Ramaekers, Enrico Amico

## Abstract

Psilocybin induces profound alterations in consciousness, yet prevailing neural models often describe a monolithic change in brain connectivity that may not fully capture the multifaceted nature of the psychedelic state. To test the hypothesis of a composite neural state, this study applied a robust, data-driven framework, Connectome Independent Component Analysis (connICA) with multi-level resampling, to resting-state fMRI data from healthy volunteers. The analysis decomposed connectomes into statistically independent functional connectivity traits ("FC-Traits"), revealing a primary trait whose expression was significantly modulated by plasma psilocin concentration, providing a whole-cortical signature of the drug’s physiological action. Crucially, a second, distinct trait was also isolated, which independently associated with impaired performance on a visual divergent thinking task. These findings demonstrate that the acute psilocybin state is a composite of co-occurring neural processes. This validates the application of a decompositional connectomic framework to move beyond global descriptions and successfully disentangle the specific neural patterns underlying distinct pharmacological and cognitive correlates.

## Introduction

The 21st century has witnessed a remarkable resurgence of scientific interest in classic psychedelic compounds, a field of inquiry that had remained largely quiescent for decades (Nichols, 2016). This "psychedelic renaissance" is propelled by a growing body of evidence demonstrating the therapeutic potential of psilocybin and other substances, which, when combined with psychological support, have shown rapid and sustained efficacy in treating a range of challenging psychiatric conditions, such as (treatment-resistant) depression (R. Carhart-Harris et al., 2021; Davis et al., 2021; Griffiths et al., 2016; Ross et al., 2016). These findings have catalysed efforts to understand the neurobiological mechanisms by which psilocybin exerts its clinical effects.

Alongside this clinical renaissance, the field of systems neuroscience has undergone its own paradigm shift. The focus has moved from mapping the function of isolated brain regions to understanding the brain as an integrated, complex system: a network of anatomically distinct but functionally coupled regions known as the functional connectome (Biswal et al., 2010). The network neuroscience (Bassett & Sporns, 2017) approach allows to explore intrinsic organization of brain dynamics into large-scale resting-state networks (RSNs), and how these support the diverse aspects of human cognition (Thomas Yeo et al., 2011). The convergence of these two fields has proven synergistic (Deco et al., 2024; Mallaroni et al., 2024; Siegel et al., 2024; Tolle et al., 2024). Indeed, psychedelic-induced perturbations of consciousness offer an unparalleled opportunity to probe the fundamental principles of whole-brain organization, while network neuroscience provides the quantitative tools to move beyond phenomenological descriptions, toward a mechanistic account of brain connectivity patterns in the psychedelic state.

From this whole-brain approach, a consistent picture of psilocybin’s acute effects on the functional connectome has emerged. The dominant finding is a large-scale reorganization of the brain’s network architecture, shifting from a state of segregated, specialized processing toward a less modular configuration (Daws et al., 2022). Recent high-precision mapping studies characterize this process as a widespread cortical desynchronization that dissolves the boundaries between functional networks (Siegel et al., 2024). This shift toward a less predictable state of brain activity, reflecting an expanded repertoire of available brain states, is a concept frequently discussed in terms of increased ’brain entropy’ (Carhart-Harris et al., 2014). At the network level, this reorganization is characterized by decreased functional connectivity within established RSNs (a process termed network "disintegration") and increased connectivity between them ("desegregation") (Carhart-Harris et al., 2012; Carhart-Harris & Friston, 2019). More recent gradient-based analyses describe this effect as a "flattening" of the brain’s principal functional hierarchy, where the typical distinction between unimodal sensory networks and transmodal association cortex is diminished (Deco et al., 2024; Girn et al., 2022). Critically, the magnitude of this brain-wide integration, measured as decreased network modularity, has been linked to the therapeutic efficacy of psilocybin (Daws et al., 2022).

This global reconfiguration is not uniform across the cortex. Previous studies have reported pronounced effects in high-level association networks, particularly the default mode network (DMN) (Gattuso et al., 2023), a system implicated in self-referential thought, introspection, and autobiographical memory (Buckner et al., 2008; Raichle, 2015). The disintegration of the DMN is a robust neural correlate of the psychedelic experience, and its magnitude correlates strongly with the intensity of subjective reports, especially the hallmark phenomenon of "ego dissolution," a transient blurring of the boundary between self and environment (R. L. Carhart-Harris et al., 2012; Lebedev et al., 2015; Smigielski et al., 2019). Furthermore, this reorganization is directly linked to the drug’s physiological action; the degree of network disintegration and increased inter-network connectivity scales directly with plasma concentrations of psilocin, the active metabolite of psilocybin (Madsen et al., 2021). Together, these findings established a foundational model linking psilocybin’s effects across physiological, neural, and phenomenological levels.

However, this DMN-centric model, while foundational, may obscure a more complex reality. The subjective psychedelic experience is not a monolithic state but a multifaceted one, encompassing profound alterations across perception, emotion, and cognition (Mason et al., 2021; Nour et al., 2016; Yu et al., 2024). It is plausible that the underlying neural state is similarly composite, consisting of multiple, co-occurring functional connectivity patterns rather than a single global reconfiguration. Uncovering such a composite structure, nevertheless, presents significant methodological challenges. Hypothesis-driven ROI analyses risk confirmation bias and can miss unexpected whole-brain patterns unless complemented by exploratory approaches. Conversely, while whole-brain analyses avoid this bias, testing every connection between brain regions entails a severe statistical penalty from multiple comparisons correction, often rendering subtle but meaningful effects undetectable (Amico et al., 2017). Thus, the field increasingly recognizes the need for data-driven approaches capable of identifying distinct whole-brain patterns or ’brain states’ associated with psychedelic action (Daws et al., 2022; Luppi et al., 2021; Singleton et al., 2022; Vohryzek et al., 2022).

To test this ’composite state’ hypothesis, we employed connectome-based Independent Component Analysis (connICA), a data-driven framework that applies ICA to the functional connectivity domain (Amico et al., 2017). While other data-driven approaches like k-means clustering are well-suited to identifying recurring temporal brain states in psychedelic data (Luppi et al., 2021), connICA is specifically designed to deconstruct a connectome into a set of statistically independent, co-occurring spatial patterns. By its mathematical nature, ICA assumes an observed signal is a mixture of independent sources; applied to the connectome, connICA is therefore well-suited to ’unmix’ the overall brain state into its constituent functional connectivity patterns, or ’FC-Traits’. This allows us to model a scenario where multiple distinct neural processes, such as a direct pharmacological effect and a separate cognitive modulation, are happening simultaneously.

For each identified FC-trait, every individual is assigned an expression weight quantifying the degree to which that pattern is present in their connectome. This decompositional approach also provides a key statistical advantage: by shifting the unit of analysis from tens of thousands of connections to the expression weights of a small number of traits, it can increase the power to detect associations with external variables like pharmacology or behaviour (Amico et al., 2017). The utility of this framework is well-established, having been successfully employed to identify clinically relevant FC-Traits in complex conditions like disorders of consciousness, alcoholism risk, and chronic cannabis use (Amico et al., 2017; Amico et al., 2020; Ramaekers et al., 2022). These prior findings demonstrate the framework’s utility and support its suitability for deconstructing the equally complex psychedelic state.

The present study, therefore, applies the connICA framework to test this ’composite state’ hypothesis directly. We aim to deconstruct the acute psilocybin-induced connectome into its constituent, independent FC-Traits to determine if the overall brain state is better described as a composite of simultaneously occurring neural processes. Specifically, we formulated a set of hypotheses to test this ’composite state’ model and establish the functional significance of any resulting data-driven patterns. We first hypothesized that the psilocybin-induced connectome could be successfully deconstructed into multiple, statistically independent FC-Traits. Our principal hypothesis was that, given prior evidence linking plasma psilocin concentration to large-scale network reorganization (Madsen et al., 2021), this decomposition would isolate at least one FC-Trait whose expression scales with the drug’s direct physiological action. This finding alone would provide evidence for a heterogeneous state comprising distinct, pharmacologically driven processes. We also exploratorily tested whether other independent traits were associated with a separate behavioural measure.

Finally, to validate the functional relevance and predictive potential of our decompositional approach, we hypothesized that the expression weights of any identified significant traits would, in combination, serve as a robust neural signature sufficient to accurately classify participants into their respective treatment groups.

## Results

### Data-Driven Discovery of Robust FC-Traits

This study first employed a data-driven analytical approach to decompose the whole-brain functional connectomes into a set of robust, independent patterns (FC-traits, see Figure 1). A multi-level stability analysis (see Methods for details) was used to empirically determine the optimal number of independent components (ICs) and identify patterns that were stable across participant subsamples.

**Figure 1.**
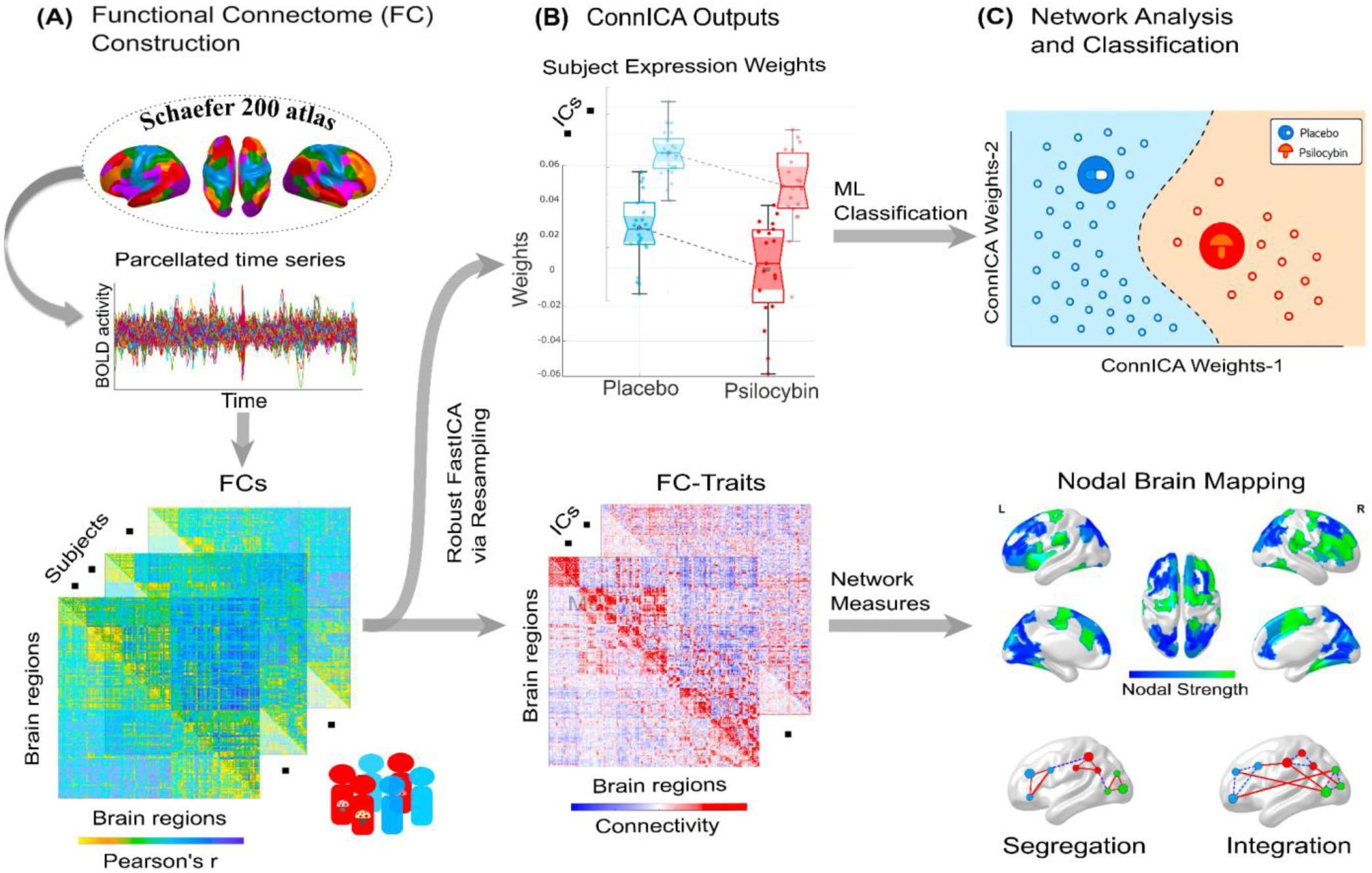
A Data-Driven Framework for Deconstructing and Analysing the Psilocybin-Induced Brain State. **(A) Functional Connectome Generation**. For each participant, mean BOLD time series were extracted from 200 cortical regions of the Schaefer atlas. Subject-specific functional connectomes (FCs) were then generated by calculating the Pearson correlation between every pair of regional time series. **(B) Connectome Decomposition**. The vectorized upper triangular elements from each participant’s symmetric FC matrix were concatenated into a single data matrix. This collection of whole-brain connectivity patterns, without prior stratification by group, was then decomposed using Robust FastICA via Resampling. This multi-level stability framework repeatedly applies the FastICA algorithm across numerous participant subsamples to identify highly reliable components. The process yields two primary outputs: FC-Traits, representing shared patterns of brain connectivity, and corresponding Subject Expression Weights, which quantify how strongly each individual expresses a given trait. **(C) Network Analysis and Classification**. The outputs were utilized for subsequent analysis and interpretation. The subject weights were used as features to train machine learning (ML) classifiers to test the predictive power of the FC-Traits in classifying group status (Psilocybin vs. Placebo). The spatial patterns of the FC-Traits were further analysed by visualizing network measures, such as nodal strength, via nodal brain mapping to identify key hubs of segregation and integration.

The analysis of model dimensionality revealed that while model fit (Adjusted R²) generally increased with the number of ICs, the rate of improvement showed a distinct inflection point around the 30-component mark (see Figure 2A). While a 30-component model was most frequently selected as optimal (in 59% of iterations), other dimensionalities were optimal for specific subsamples, reinforcing the need for our flexible, data-driven approach (see Figure 2B).

**Figure 2.**
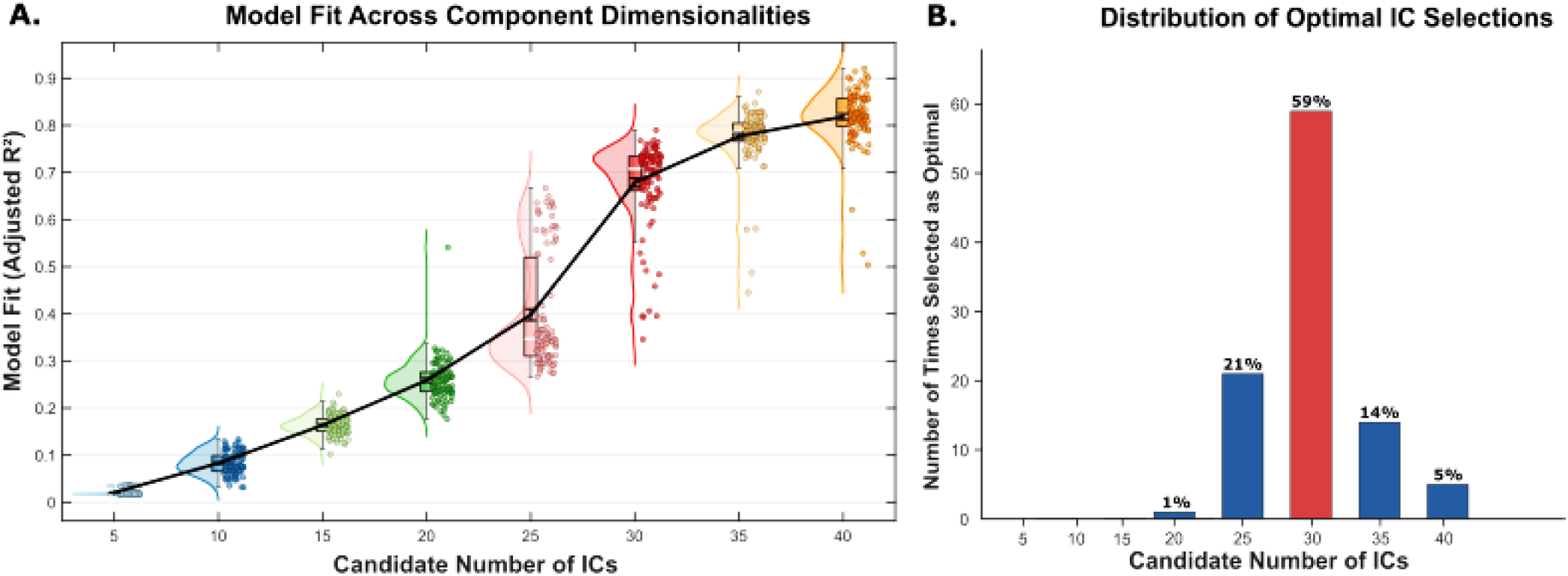
Data-Driven Selection of the Optimal Number of Independent Components. The figure displays results from the 100-iteration resampling analysis used to determine the optimal number of independent components (ICs) for the connICA decomposition. **(A)** The raincloud plots show the distributions of model fit (Adjusted R²) for each of the eight tested numbers of ICs. For each number of ICs, the corresponding plot shows the distribution of Adjusted R² values from the 100 resampling iterations, where each data point represents the outcome of a single iteration. This visualization reveals that while the mean model fit generally increases with the number of ICs, the rate of improvement plateaus around 30 ICs. **(B)** The bar chart illustrates the frequency with which each number of ICs was selected as optimal across the 100 iterations. The optimal number of ICs for each iteration was identified by maximizing the variance explained per robust component (Adjusted R² / Number of RCs).

Robust components derived from applying ICA with the best model order (i.e., number of ICs) on 100 subsamples were aggregated, yielding a total of 2,334 components for a final cross-iteration stability analysis. From this collection, hierarchical clustering and a 60% robustness threshold were used to identify consistently expressed patterns. This procedure yielded a final set of 21 robust FC-traits (see Supplementary Table 1 for robustness scores of the stable clusters).

### Functional Significance of FC-Traits

To ascertain their functional significance, the 21 robust FC-traits were tested for associations with pharmacological, behavioural, and subjective variables (see Methods). After applying a False Discovery Rate (FDR) correction for multiple comparisons, this analysis revealed two distinct and functionally relevant traits: a primary trait whose expression was directly modulated by plasma psilocin concentration, and a second trait associated with a measure of divergent thinking.

### The Psilocin-Associated Functional Trait (PA-FT)

The first brain pattern discovered, termed the Psilocin-Associated Functional Trait (PA-FT), is detailed in Figure 3. The expression of this trait was significantly suppressed in the psilocybin group relative to the placebo group (see Figure 3A). The multiple linear regression model (*R*^2^= 0.33) confirmed that plasma psilocin concentration measured at 80 minutes (the observed peak) was a highly significant predictor of the PA-FT weights (FDR-corrected p = 0.0348), with higher physiological drug levels corresponding to a weaker (more negative) expression of this trait (see Figure 3B). This trait was also found to be significantly associated with subjective ratings on the Ego Dissolution Inventory (EDI); further details on this secondary association are provided in the Supplementary Materials (see Supplementary Figure 2).

**Figure 3.**
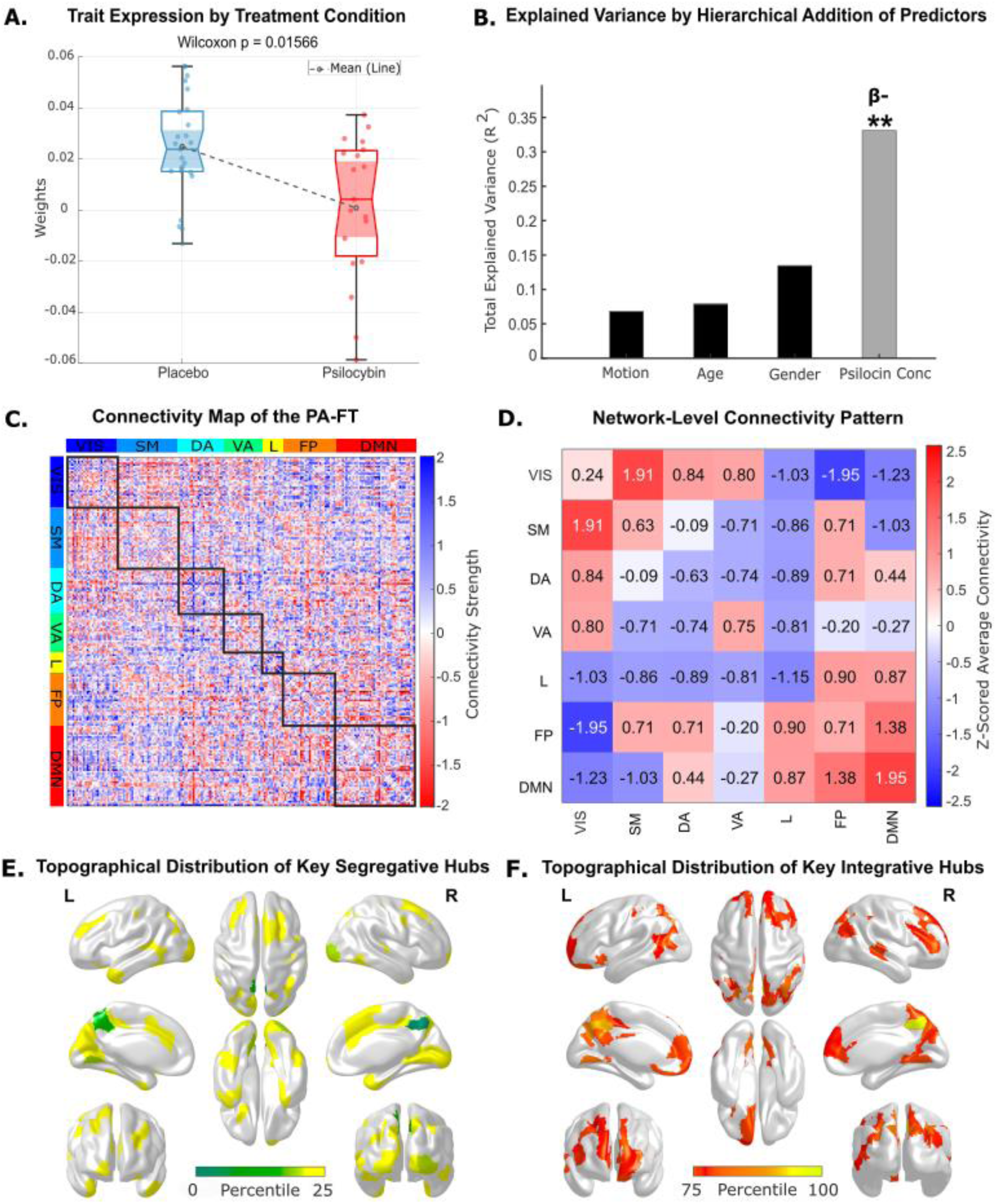
Characterization of the Psilocin-Associated Functional Trait (PA-FT). **(A)** The box plot displays the distribution of subject-specific weights for the PA-FT, separated by treatment group. The expression of this trait is significantly suppressed in the psilocin group relative to the placebo group (Wilcoxon p = 0.0157). **(B)** The bar chart illustrates the incremental R-squared from the multiple linear regression model predicting PA-FT weights. After accounting for covariates, plasma psilocin concentration emerges as a highly significant predictor, with the full model explaining approximately 33% of the variance. The asterisks (**) denote the statistical significance of the psilocin concentration’s contribution to the model (uncorrected p < 0.005), while the negative beta sign (β-) indicates that higher psilocin levels are associated with weaker expression of this trait. **(C)** The full 200*200 spatial map of the PA-FT, with nodes ordered according to the seven canonical resting-state networks. Red values indicate positive connectivity, while blue values denote negative connectivity. **(D)** A 7*7 summary matrix showing the Z-scored average connectivity within each network (on the diagonal) and between all pairs of networks (off-diagonal). The matrix reveals a dominant pattern of positive connectivity between early sensory networks (Visual, Somatomotor) and within the DMN. **(E-F)** Surface projections display the topographical distribution of the key cortical hubs defining the trait, based on nodal strength. The maps show the top 25% of brain regions, ranked separately by their **(E)** negative (segregative) and **(F)** positive (integrative) nodal strength.

The PA-FT’s connectivity profile is simultaneously characterized by strong positive and negative couplings. The prominent negative couplings primarily link the Visual network with higher-order systems, while strong positive associations are observed between the Visual and Somatomotor networks, between the Default Mode (DMN) and Fronto-parietal (FP) networks, and notably, within the DMN itself (see Figures 3C-D).

The PA-FT’s nodal strength distribution highlights the Visual and Default Mode networks as architecturally central. These systems hosted the majority of hubs with the strongest negative strength (segregative hubs) and, alongside the FP network, also contained hubs with the strongest positive strength (integrative hubs). This co-location of opposing hub types within the same large-scale network points to a functional heterogeneity, where distinct subregions drive different connectivity patterns (see Figures 3E-F).

### Visual Divergent Thinking-Associated Functional Trait (VDT-FT)

The second significant brain pattern, which we termed the Visual Divergent Thinking-Associated Functional Trait (VDT-FT), is fully characterized in Figure 4. The expression of this trait was significantly lower in the psilocybin group compared to the placebo group (see Figure 4A). Subsequent multiple linear regression analysis confirmed that divergent creative performance on the Picture Concept Task (PCT) was a highly significant predictor of the VDT-FT weights. Specifically, PCT Fluency was a highly significant predictor (*R*^2^ = 0.366, FDR-corrected p = 0.0069), with weaker expression of this specific connectivity pattern corresponding to impaired creative performance (see Figure 4B). The PCT Originality score was also found to be significantly associated with this trait; however, as these two measures were highly correlated (Pearson’s r ≈ 0.78), we present this association in the Supplementary Materials (see Supplementary Figure 3) to avoid collinearity.

**Figure 4.**
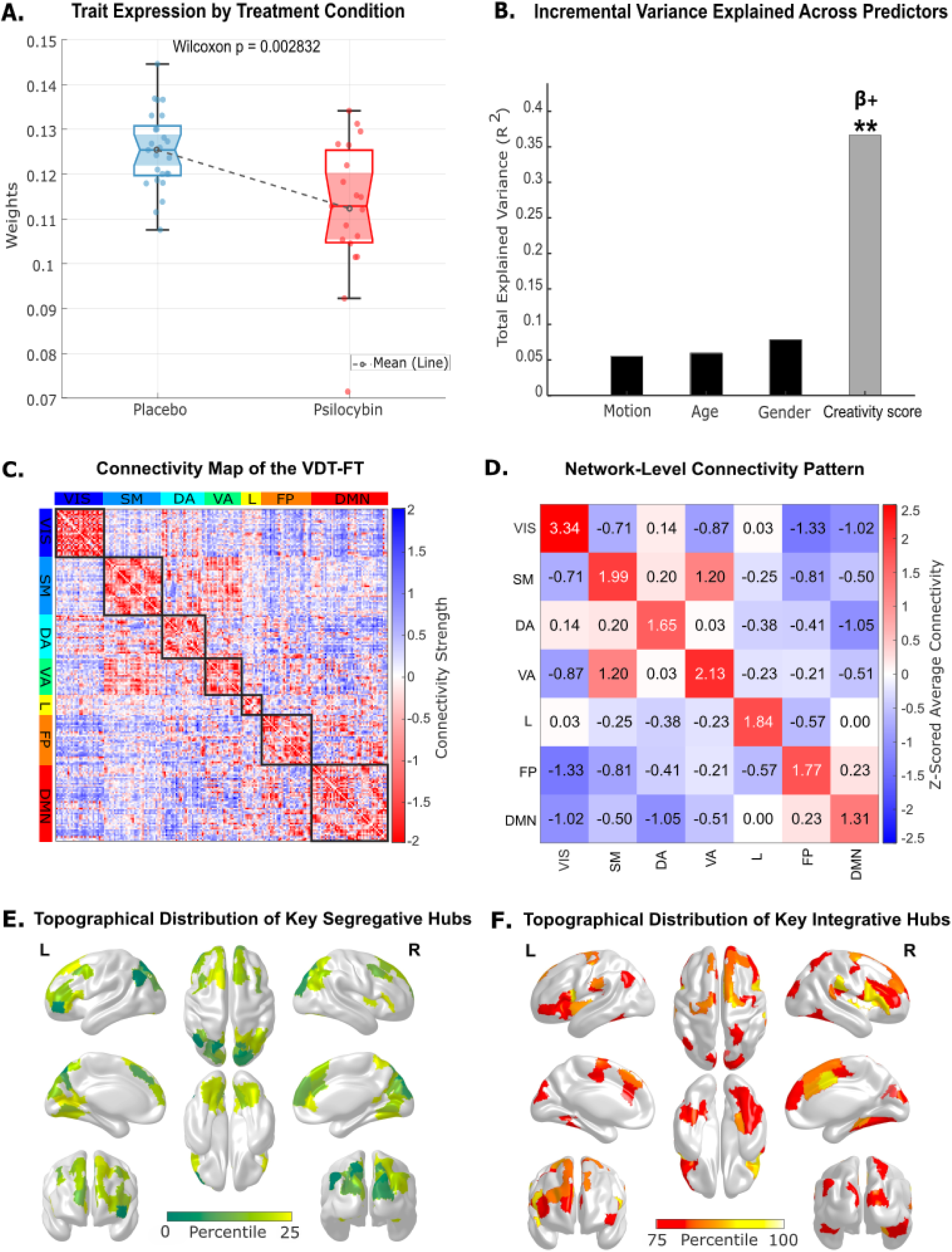
Characterization of the Visual Divergent Thinking-Associated Functional Trait (VDT-FT). **(A)** The box plot displays the distribution of subject-specific weights for the VDT-FT, separated by treatment group. The expression of this trait is significantly lower in the psilocybin group relative to the placebo group (Wilcoxon p = 0.0028). **(B)** The bar chart illustrates the incremental R-squared from the multiple linear regression model predicting VDT-FT weights. After accounting for covariates, the Picture Concept Task (PCT) fluency score emerges as a highly significant predictor, with the full model explaining approximately 36.6% of the variance. The asterisks (**) denote the statistical significance of the creativity score’s contribution to the model (uncorrected p < 0.005), while the positive beta sign (β+) indicates that higher creative fluency is associated with stronger expression of this trait. **(C)** The full 200*200 spatial map of the VDT-FT, with nodes ordered according to the seven canonical resting-state networks (Thomas Yeo et al., 2011). Red values indicate positive connectivity, while blue values denote negative connectivity. **(D)** A 7*7 summary matrix showing the Z-scored average connectivity within each network (on the diagonal) and between all pairs of networks (off-diagonal). The dominant pattern is positive within-network connectivity. **(E-F)** Brain renders of the top 25% of brain regions ranked by their nodal strength, illustrating the topographical distribution of the key hubs that define the trait. The maps are separated into **(E)** negative hubs, which drive the trait’s segregative properties, and **(F)** positive hubs, which drive its integrative properties.

The spatial architecture of the VDT-FT, shown in detail in Figure 4C, is defined by a pattern of strong functional integration mainly within early sensory and attentional networks, coupled with segregation between the visual network and higher-order association cortices. The network-level summary in Figure 4D clarifies this signature, showing strong positive connectivity within the Visual (VIS), Somatomotor (SM), and Ventral Attention (VA) networks, alongside negative connectivity between the VIS network and both the Frontoparietal (FP) and Default Mode (DMN) networks.

Analysis of the VDT-FT’s nodal strength distribution revealed that its strongest segregative hubs were primarily located in the Visual and Dorsal Attention networks (see Figure 4E). The main integrative hubs were also concentrated in these same systems, along with the Somatomotor network (see Figure 4F). Together, this distribution suggests that the VDT-FT is shaped primarily by a core set of sensory-motor and attentional circuits, which jointly contribute to both the integration and segregation underlying this creativity-related brain state.

Beyond the two significant findings reported above, no other associations between any of the 21 robust FC-traits and the remaining acute creativity measures or subjective measures survived FDR correction.

### Classification Performance of Combined FC-Traits

To determine if the identified FC-traits could serve as a robust neuroimaging-informed signature, we evaluated their combined ability to classify participants into psilocybin or placebo groups solely based on their connICA patterns. The subject-specific weights of the VDT-FT and PA-FT were used as features to train three machine learning models, with performance evaluated using a 10-fold cross-validation procedure (see Figure 5).

**Figure 5.**
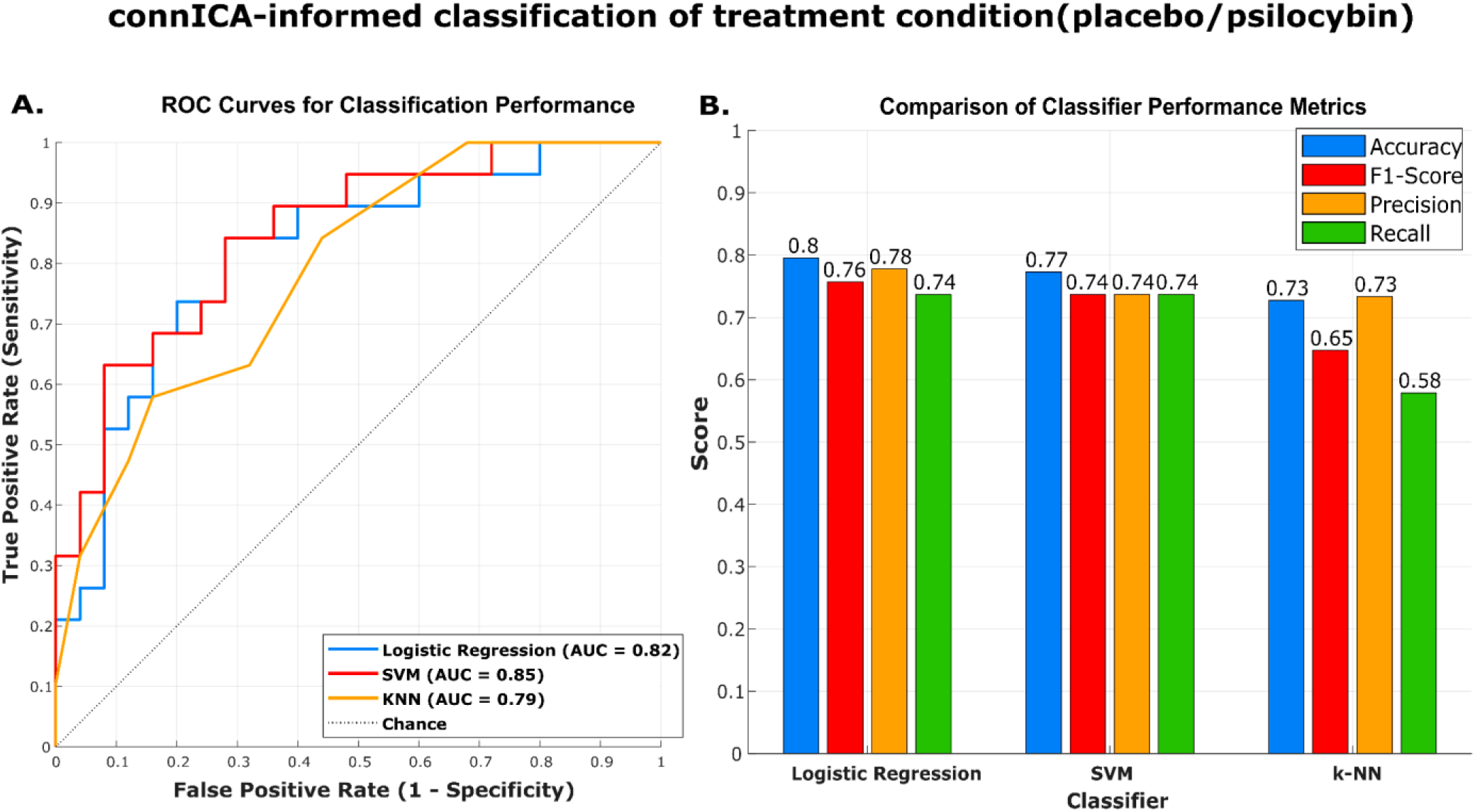
Cross-Validated Classification Performance of the Combined FC-Traits. This figure displays the performance of three machine learning models trained to classify participants into psilocybin or placebo groups using the combined FC-traits as features. **(A)** Receiver Operating Characteristic (ROC) curves show the classifier performance across all decision thresholds. The Area Under the Curve (AUC) was highest for the SVM (0.85), followed by Logistic Regression (0.82) and k-NN (0.79). The dotted line represents chance performance. **(B)** Bar plot comparing key performance metrics. Logistic Regression achieved the best overall balance of metrics with an accuracy of 0.80, F1-Score of 0.76, precision of 0.78, and recall of 0.74. Together, these results show that the combined FC-traits can reliably distinguish between the two treatment states.

The linear classifiers yielded the strongest results. While the Support Vector Machine (SVM) showed the highest discriminative capacity (AUC = 0.85), the Logistic Regression model achieved the highest classification accuracy (80%) with a comparable AUC (0.82). The robust performance of these linear models, which surpassed the non-linear k-NN classifier, suggests that the distinction between treatment groups is encoded in a linearly separable manner within the two FC-traits. This level of classification performance demonstrates that the combined FC-traits can reliably distinguish an individual’s condition (psilocybin or placebo) from their functional connectomes.

## Discussion

This study employed a data-driven connectome decomposition framework to investigate the constituent neural patterns underlying the acute psilocybin state. The principal finding is that the complex, whole-brain reorganization induced by psilocybin can be deconstructed into statistically independent and functionally distinct components. Crucially, our analysis isolated a Psilocin-Associated Functional Trait (PA-FT) whose expression was modulated by plasma psilocin concentration, providing a robust neural signature of the drug’s physiological action. Independently, we identified a second component, a Visual Divergent Thinking-Associated Functional Trait (VDT-FT), whose expression was associated with behavioural measures of creative cognition.

This decomposition offers a more granular perspective on the prevailing models of the psychedelic state, which is characterized by global shifts in brain dynamics such as increased entropy or a uniform collapse of network segregation (R. L. Carhart-Harris et al., 2014; Madsen et al., 2021). While such global descriptions have been foundational, the rich and varied phenomenology of the psychedelic experience, encompassing changes in perception, self-consciousness, and higher-order cognition, suggests a more complex underlying neural architecture (Mason et al., 2021; Nour et al., 2016). Our findings provide empirical support for this view by demonstrating that the psilocybin-induced state is a composite of distinct neural processes. The successful identification of the PA-FT serves to validate the connICA framework’s ability to detect a robust, physiological trait of psilocybin’s action. The distinct value of this approach is then demonstrated by its capacity to isolate, on top of this global drug-induced pattern, an independent functional trait (the VDT-FT) that predicts a behavioural outcome. This methodological advance suggests the potential for this framework to be used to identify other clinically relevant FC-traits, such as a neural marker capable of predicting therapeutic response. More broadly, this decompositional approach itself offers a conceptual advance, providing a framework where the interplay between different neural patterns, modulated to varying degrees by the drug, could potentially offer a mechanistic basis for understanding inter-individual variability in psychedelic effects (Mallaroni et al., 2024b; Tolle et al., 2024), moving beyond a single, ’average’ state description.

### A Plasma Concentration-Dependent Brain Signature of Psilocybin Action

The primary significant component identified, the PA-FT (See Figure 3), represents a data-driven pattern reflecting key aspects of psilocybin’s physiological effects on whole-brain functional connectivity. The expression of this trait was significantly suppressed in the psilocybin group relative to placebo, and critically, the degree of suppression correlated significantly with plasma concentrations of psilocin. This finding provides empirical support for a growing body of work demonstrating concentration-dependent effects of psychedelics on brain network organization (Madsen et al., 2021; Olsen et al., 2022).

Despite evidence that psilocybin alters activity across multiple brain networks, the field has emphasized the role of the DMN (R. L. Carhart-Harris et al., 2012; Smigielski et al., 2019). The spatial map of the PA-FT includes strong positive within-DMN connectivity (see Figure 3D). Therefore, the observed suppression of this trait (i.e., lower individual weights under psilocybin corresponding to higher psilocin levels) aligns well with prior findings, specifically the work by Madsen et al. (2021) which demonstrated that higher plasma psilocin concentrations correlate with reduced DMN integrity (coherence) and decreased overall network segregation.

However, the data-driven nature of our approach allows us to characterize this concentration-dependent effect beyond pre-hypothesized regions of interest. While Madsen et al. (2021) showed correlations between psilocin levels and global metrics of integrity and segregation, and Olsen et al. (2022) linked psilocin levels to changes in the occurrence of specific dynamic connectivity states, our connICA method isolates a stable, static FC pattern (the PA-FT) whose overall expression is significantly associated with plasma psilocin. This data-driven pattern confirms that the concentration-dependent effect is not confined to the DMN; indeed, the PA-FT also encompasses strong negative connectivity between the Visual network and higher-order DMN and FP networks, alongside positive connectivity between Visual and Somatomotor systems (see Figures 3C-D). This broader, multi-network pattern resonates with recent perspectives emphasizing psychedelics’ widespread impact across the cortex and aligns with findings suggesting large-scale cortical desynchronization as a key feature of the acute state (Siegel et al., 2024). The PA-FT can thus be interpreted as a specific, identifiable signature reflecting this broader concentration-dependent network reorganization. This network reorganization has been previously linked to the intensity of ’ego dissolution’ (Tagliazucchi et al., 2016), providing a rationale for the secondary association we also found between the PA-FT’s expression and subjective EDI scores (see Supplementary Figure 2). The distribution of the trait’s key hubs, prominent in both the DMN and Visual networks (see Figures 3E-F), reinforces that psilocybin’s effects linked to plasma concentration manifest across multiple interacting brain systems.

### A Whole-Brain Substrate for the Suppression of Visual Divergent Thinking

The second significant component identified, the VDT-FT (See Figure 4), was linked to cognitive performance. Specifically, its expression was positively correlated with divergent thinking performance on the Picture Concept Task (PCT). The expression of this trait was also significantly suppressed in the psilocybin group. This finding provides a compelling neural substrate for the nuanced effects of psilocybin on creativity reported by Mason et al. (2021). Specifically, our results suggest that the VDT-FT represents the functional connectome substrate of this deliberate form of creative cognition, and its suppression under psilocybin explains the observed performance decrements.

While Mason et al. (2021) linked this behavioural impairment to altered connectivity between specific networks like the DMN and FPN, our data-driven approach identifies a holistic, whole-brain functional trait whose expression is associated with performance on this specific task. The spatial architecture of the VDT-FT is defined by strong integration within primary sensory and attentional networks, which are simultaneously segregated from transmodal association cortices like the DMN and FPN (see Figures 4C-D). This configuration appears organized for externally cued tasks that require focused attention on sensory input while minimizing interference from internal thought streams (Beaty et al., 2016).

By reducing the expression of the VDT-FT, psilocybin impairs this cognitive architecture. A plausible mechanism for this disruption lies in psilocybin’s primary action on transmodal cortices. Association cortices, such as the prefrontal cortex, are known to be enriched with 5-HT_2A_ receptors, particularly on deep-layer pyramidal neurons (Kwan et al., 2022). Psilocybin’s agonism of these receptors increases neuronal excitability, which likely disrupts the coordinated activity required to maintain segregation between these higher-order systems and the sensory (visual) cortices engaged by the task. The lower trait weights under psilocybin therefore signify a breakdown of this crucial decoupling, explaining the observed performance decrements. The nodal strength analysis, which localizes the trait’s most influential hubs to these same sensory and attentional systems (see Figures 4E-F), further corroborates this functional specialization.

It is notable that the VDT-FT’s expression was not associated with divergent creative performance on the AUT. This specificity may be a meaningful finding: the VDT-FT’s architecture is dominated by visual and attentional networks, aligning well with the visual-associative demands of the PCT. The AUT, in contrast, is primarily a verbal task reliant on semantic memory retrieval. This suggests the VDT-FT captures a neural substrate specific to visual-perceptual divergent thinking, while semantic-based divergent thinking likely depends on a different neural architecture not captured by this specific trait.

### Methodological Advances and Predictive Power of Functional Connectome Traits

The findings of this study were enabled by a methodological framework that overcomes several limitations of traditional neuroimaging analyses. By employing connICA, we shifted the unit of analysis from tens of thousands of individual connections to a small number of whole-brain connectivity patterns, thereby circumventing the severe statistical burden of mass-univariate testing and increasing the power to detect functionally relevant associations (Amico et al., 2017). Furthermore, the novel multi-level resampling and stability analysis ensured that the identified FC-Traits were highly robust and not artifacts of a specific participant subsample or an arbitrary choice of the number of ICs.

A key validation of this approach was given by the predictive power of the identified FC-traits. In fact, the subject-specific weights of the VDT-FT and PA-FT were sufficient to classify participants into their respective treatment groups with high accuracy (80% accuracy, AUC = 0.85, See Figure 5). It confirms that our framework successfully distilled the complex, high-dimensional psychedelic state into a potent and low-dimensional neural signature, establishing that the subject-specific weights derived from these functional connectome traits provide a validated and meaningful basis for accurately distinguishing individuals in the acute psilocybin state from those under placebo.

### Limitations and Future Directions

While this study provides novel insights, it is important to consider its limitations, which suggest directions for future research. First, the findings are based on a cohort of healthy volunteers with prior psychedelic experience. To establish the robustness and generalizability of the discovered FC-traits, replication is needed in larger, more diverse cohorts, including psychedelic-naive individuals and clinical populations, such as patients with depression, for whom psilocybin shows therapeutic promise. Additionally, the parallel-group design used here is effective for group-level comparisons but cannot resolve how these FC-trait weights change within an individual relative to their own baseline. Future work would benefit from a within-subject, crossover design to provide a more powerful assessment of individual-level modulation of these connectomic traits.

Second, the present analysis focused on characterizing connectomic patterns at the cortical level. A future direction will be to extend this decompositional framework to include subcortical structures. The thalamus, for instance, is a central hub rich in 5-HT_2A_ receptors and is integral to the "thalamic gating" hypothesis, which suggests psychedelics disrupt the filtering of sensory information to the cortex (Geyer & Vollenweider, 2008). Moreover, preserved thalamocortical connectivity has been shown to distinguish the psychedelic state from states of reduced consciousness (R. L. Carhart-Harris et al., 2013). Applying a whole-brain (cortical and subcortical) connICA approach is therefore necessary to fully characterize the role of these critical cortico-striato-thalamo-cortical (CSTC) loops in the psychedelic state.

Building on the current work, future research with larger cohorts could leverage these FC-trait weights as features in connectome-based predictive models to forecast continuous clinical or behavioural outcomes, moving beyond classification to predict the intensity of subjective effects or long-term therapeutic response. Furthermore, our analysis only tested associations against plasma concentration and creativity; the other robust FC-traits identified by our decomposition remain to be characterized. These traits should be explored in future studies against a wider battery of subjective, clinical, and cognitive variables to determine their full functional relevance. Additionally, a promising next step would be to apply modularity analysis to the identified FC-Trait matrices. Modularity provides a quantitative measure of network segregation, the degree to which a network can be subdivided into distinct, densely intra-connected communities (Newman, 2006). This metric would offer a direct, graph-theoretical test of the widespread hypothesis that the psychedelic state reflects a shift from a segregated to a more integrated brain architecture (R. L. Carhart-Harris et al., 2014). It would be predicted that the PA-FT, representing the baseline state of brain organization, would exhibit a significantly higher modularity score than the connectome of an individual under the influence of the drug.

### Conclusion

In conclusion, this study demonstrates that the acute psilocybin-induced brain state is not a monolithic neural event but a composite of distinct, co-occurring functional patterns. By employing a robust, data-driven decompositional framework, we successfully isolated a primary neural signature of the drug’s physiological action, which scales with plasma psilocin concentration, from a separate, independent pattern linked to performance on a visual divergent thinking task. This work validates an analytical approach for psychedelic neuroscience, establishing a framework to disentangle the constituent neural substrates of multifaceted neurocognitive states and map them to their specific pharmacological and cognitive correlates.

## Methods

### Participants

This study employed data from a clinical trial conducted at Maastricht University, which employed a randomized, double-blind, placebo-controlled, parallel-group design. The trial recruited 60 healthy volunteers who were matched for age, sex, and education and had prior psychedelic experience, though not within the three months preceding the study. Participants received either 0.17 mg/kg psilocybin or a placebo. All procedures were ethically approved in accordance with the Declaration of Helsinki, and all participants provided written informed consent. Full procedural details are available in the primary publications (Mason et al., 2020, 2021).

Of the 60 participants recruited, 49 provided resting-state fMRI (rs-fMRI) data, from which a final sample of 44 was selected for the present analysis following a quality control procedure. Exclusions were made due to unavailable plasma psilocin data (n=3, one of whom also exhibited >25% motion artifacts), excessive head motion (>10% artifact volumes, n=1), and missing BOLD data across five cortical parcels (n=1). Conversely, one participant with missing BOLD data from a single parcel was retained after the data were imputed. The final cohort therefore comprised 19 individuals in the psilocybin group and 25 in the placebo group.

### Image Acquisition

All neuroimaging data were acquired on a MAGNETOM 7T MRI scanner. The scanning session included a high-resolution T1-weighted structural scan at approximately 50 minutes post-administration for anatomical reference and registration. This was followed by a 6-minute rs-fMRI scan at 102 minutes post-administration, timed to capture brain activity near the peak plasma concentration (observed at 80 minutes post-administration).

The rs-fMRI data were collected using a gradient-echo echo-planar imaging (EPI) sequence, during which 258 whole-brain volumes were acquired. The acquisition parameters were as follows: TR=1400 ms, TE=21 ms, flip angle=60°, FOV=198 mm, 72 interleaved slices, and 1.5 mm isotropic voxels. Throughout the acquisition, participants were instructed to keep their eyes open, focus on a central crosshair, remain still, and try to stay awake while clearing their mind (Mason et al., 2020).

### fMRI Data Preprocessing and Parcellation

Resting-state fMRI data were pre-processed using the Apéro MATLAB toolbox (Mallaroni et al., 2024; Tolle et al., 2024), which implements the connectome pipeline of (Amico et al., 2017) using FSL routines (Jenkinson et al., 2012).

High-resolution T1-weighted images were denoised, skull-stripped (HD-BET), segmented into gray matter (GM), white matter (WM), and CSF, and non-linearly registered to MNI-152 space (Tolle et al., 2024). The Schaefer 200-parcel, 7-network atlas (Schaefer et al., 2018) was selected to define network nodes. This atlas was chosen for its demonstrated capacity to produce connectomes with high topological stability across different preprocessing pipelines, making findings less susceptible to methodological artifacts (Luppi & Stamatakis, 2021). The volumetric atlas was warped into each participant’s native space using nearest-neighbour interpolation to preserve parcel integrity (Tolle et al., 2024).

The functional data processing pipeline included slice-timing correction, skull-stripping (FSL BET), motion correction (FSL MCFLIRT), intensity normalization, and linear detrending. A multi-stage denoising procedure was applied prior to filtering, which involved an 18-parameter nuisance regression (including motion, tissue signals, global signal, and their derivatives) followed by a component-based correction where the first three principal components were regressed out (Tolle et al., 2024). The cleaned, unsmoothed time series were then band-pass filtered (0.01-0.25 Hz) (Tolle et al., 2024). To ensure accurate alignment for time series extraction, the tissue masks and GM parcellation derived from the T1-weighted image were registered to each subject’s mean functional volume using a 6-df rigid-body transformation followed by boundary-based registration (BBR) (Tolle et al., 2024). Finally, a single mean time series was extracted for each of the 200 cortical parcels by averaging the fully pre-processed voxel-wise data within each region.

### Head Motion Assessment and Covariate Generation

To account for potential confounds from in-scanner head motion, a single summary metric was computed for each participant and included as a covariate in all subsequent statistical analyses. This was particularly important given that the psilocybin group exhibited significantly more head motion than the placebo group (Tolle et al., 2024). Motion-contaminated volumes were identified using a multi-metric approach. Specifically, a volume was flagged as an artifact if it exceeded established thresholds on any of three distinct metrics: Frame Displacement, DVARS, or the standard deviation of the BOLD signal (Tolle et al., 2024). A comprehensive binary outlier vector was generated for each participant using a logical OR operation across these three criteria. The total number of flagged volumes was then divided by the total scan length to yield a final ’motion outlier fraction’, which represents the proportion of motion-corrupted data for each subject.

### Pharmacokinetic, Behavioural, and Subjective Measures

In addition to neuroimaging data, several other measures were collected to characterize the pharmacological, cognitive, and subjective effects of psilocybin.

#### Pharmacokinetic Data

To quantify drug exposure, venous blood samples were collected at 80-, 150-, and 360-minutes post-administration, yielding mean (±SD) psilocin concentrations of 15.61 (±7.24) ng/mL, 12.86 (±4.93) ng/mL, and 4.85 (±2.35) ng/mL, respectively. These samples were analysed using liquid chromatography-mass spectrometry (LC-MS/MS) to determine plasma concentrations of psilocin, the primary psychoactive metabolite of psilocybin (Mason et al., 2020). For the purposes of our analysis, the psilocin concentration measured at the time point closest to the rs-fMRI scan was used as a key predictor variable.

#### Behavioural Measures of Creativity

Creative cognition was assessed during the acute drug effects (∼120-130 minutes post-dose) using two established tasks (Mason et al., 2021). These tasks were selected to probe the two primary components of creative thought: divergent thinking, defined as the process of generating multiple unique ideas from a single prompt, and convergent thinking, which involves identifying a single optimal solution to a well-defined problem (Guilford, 1967). The Picture Concept Task (PCT) was administered to measure both convergent (number of correct solutions) and divergent (fluency and originality of alternative ideas) thinking. In this task, participants were presented with visual stimuli (a matrix of color pictures) and asked to find associations between them. The Alternate Uses Task (AUT) served as a dedicated measure of divergent thinking, requiring participants to list as many uses as possible for common household items presented as textual prompts, providing metrics of fluency, and the generation of novel ideas, (Mason et al., 2021). These five specific acute scores (PCT-Convergent, PCT-Fluency, PCT-Originality, AUT-Fluency, and AUT-Novelty) constituted the set of behavioural variables of interest used in the regression analyses.

#### Subjective Measures

To characterize the acute psychedelic experience, subjective effects were assessed retrospectively at 360 minutes post-administration using two established questionnaires (Mason et al., 2020). These included the 5 Dimensions of Altered States of Consciousness (5D-ASC) scale (Dittrich, 1998; Studerus et al., 2010), which assesses a range of subjective experiences including perceptual and emotional changes, and the Ego Dissolution Inventory (EDI) (Nour et al., 2016). The specific subscale scores derived from these questionnaires were included in the set of variables of interest tested in the subsequent regression analyses.

### Functional Connectome Generation

To investigate the brain’s large-scale network architecture, a 200*200 subject-specific functional connectome (FC) was generated for each participant. This network representation models anatomically distinct regions (nodes) and the statistical dependencies between their activity (edges) to enable the quantitative analysis of intrinsic brain dynamics (Bullmore & Sporns, 2009). The nodes were defined by the 200 cortical parcels of the Schaefer atlas (Schaefer et al., 2018). The weight of each edge, representing the functional connectivity between two nodes, was defined by the Pearson correlation coefficient between the time series of their spontaneous BOLD signal fluctuations (Fox & Raichle, 2007; see Figure 1A).

### Connectome Independent Component Analysis (connICA)

To deconstruct the complex, whole-brain connectivity patterns into distinct, functionally relevant components, we employed connICA, a data-driven framework for analysing collections of individual functional connectomes (Amico et al., 2017). The core of this method is the application of Independent Component Analysis (ICA) to a single data matrix where each row represents the vectorized upper-triangular elements of a single participant’s symmetric FC matrix. This decomposition yields two primary outputs: a set of independent components, termed FC-Traits, which represent shared, system-level patterns of functional connectivity across the cohort; and a corresponding matrix of subject weights, which quantifies the degree to which each individual participant expresses each FC-Trait (see Figure 1B; see Supplementary Figure 1 for the full workflow).

A primary challenge in applying ICA is that the optimal number of components (i.e., the model order) is unknown a priori. Furthermore, the FastICA algorithm is non-deterministic, meaning that its solutions can vary across different runs due to random initializations (Hyvärinen & Oja, 2000). To address these challenges and ensure our findings were highly robust and not biased by the specific composition of our sample, we developed a novel, five-step resampling and stability analysis to systematically identify robust FC-Traits by testing their stability across different participant subsets and a range of model orders (by varying the number of ICs).

#### Step 1: Stratified Subsampling for Robustness Testing

The analysis began with a stratified subsampling procedure to assess the stability of FC-Traits across different participant cohorts. Over 100 iterations, we subsampled 40-subject datasets (out of the original dataset of 44 subjects) by randomly excluding two participants from the psilocybin group and two from the placebo group, thereby preserving the original balance between groups in each iteration.

#### Step 2: Robust Component Extraction within Subsamples

For each of the 100 subsampled datasets, we performed a two-layered stability analysis to identify an optimal and robust set of components. To systematically explore model dimensionality, we tested eight number of independent components (ICs) choices, increasing (in increments of 5) from 5 ICs up to the theoretical maximum of 40 ICs, a limit constrained by the number of subjects (n=40) in each subsampled data matrix. Secondly, to mitigate ICA’s non-deterministic solutions, each of these eight choices was itself repeated 100 times. Finally, we identified a set of robust components (RCs) for each ICs choice, defined as components that were highly consistent (i.e., FC-trait and weight similarity or Pearson’s r > 0.75) across more than 70% of the runs.

#### Step 3: Optimal Model Selection

For each subsampling iteration, an optimal number of ICs was selected from the eight tested choices. This was achieved by identifying the set of RCs that maximized the variance explained per number of ICs (i.e., Adjusted R² / Number of RCs), a metric that balances model fit with parsimony (see Supplementary Note for the reconstruction-based regression procedure used to calculate this Adjusted R² value). This procedure was repeated for all 100 subsampled datasets, yielding 100 sets of optimal RCs.

#### Step 4: Identification of Stable FC-Traits via Cross-Iteration Clustering

All 100 sets of optimal RCs were aggregated for the final “discovery” stage. To identify the most consistently expressed FC-Traits, we performed hierarchical agglomerative clustering on this collection based on spatial similarity (Himberg et al., 2004). To account for ICA’s inherent sign ambiguity, the absolute Pearson correlation was used as the similarity metric (|r| ≥ 0.70), grouping components by their underlying spatial pattern irrespective of polarity; subject weights were not used, as they are not comparable across different subsamples. Clusters were defined as stable, data-driven FC-Traits if their robustness score exceeded 60%, meaning they contained highly similar components in at least 60 of the 100 iterations. For each stable trait, a final representative spatial map, or centroid vector, was calculated by averaging the sign-aligned spatial vectors of all its member components.

#### Step 5: Derivation of Final Subject Weights for Stable Traits

The final step involved deriving a single, stable weight for each of the original 44 participants for every “final” FC-Trait, which is represented by the centroid vector of a robust component cluster. For a given trait, we first identified all the individual components belonging to its cluster from across the resampling iterations. The original subject weight vector for each of these member components was then retrieved. A critical sign-correction was applied to ensure all components were directionally aligned with the trait’s centroid; if a component’s spatial map was negatively correlated with the centroid, the sign of its entire subject weight vector was inverted (as ICA is blind to polarity). A participant’s final weight for the trait was then calculated by averaging all their available, sign-corrected weights from the components within that cluster. A participant’s weight from a given component was only available for this averaging procedure if they were part of the 40-subject subsample from which that component was originally derived. These final, robust subject weights served as the dependent variables in all subsequent statistical analyses.

### Network Analysis: Nodal Strength and Hub Identification

To identify the most influential brain regions (hubs) within each stable FC-Trait, we employed nodal strength, a measure that quantifies the overall connectivity of a given region (node). For each of the 200 cortical nodes in a given FC-Trait matrix, we first calculated two raw strength metrics. Positive nodal strength was computed as the sum of all positive connection weights associated with that node. Negative nodal strength was computed as the direct sum of all negative connection weights.

To isolate the most critical hubs for visualization and interpretation, we then thresholded the resulting distributions of these 200 nodal strength values (see Figure 1C). Primary integrative hubs were defined as nodes with positive strength values in the 75th-100th percentile. Primary segregative hubs were defined as nodes with negative strength values in the 0-25th percentile, such that more negative values indicate greater segregative influence (Rubinov & Sporns, 2010).

### Regression Analysis of FC-Trait Associations

To assess the functional significance of the identified FC-Traits, a series of multiple linear regression analyses was performed. For each stable FC-Trait, a separate linear model was constructed to test its association with each variable of interest individually (e.g., plasma psilocin concentration or a specific creativity score).

In every model, the dependent variable was the final, stable subject-specific weight vector for that trait. The model’s independent variables consisted of the single primary predictor of interest, along with three covariates of no interest: participant age, sex, and the "motion outlier fraction" summary statistic, as previously described. The full model took the general form:

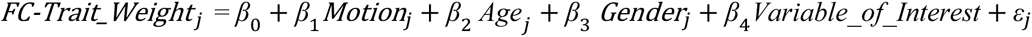

To account for multiple statistical tests (one for each robust FC-Trait against each variable of interest), we controlled the false discovery rate using the Benjamini–Hochberg procedure (Benjamini & Hochberg, 1995). Associations with FDR-corrected p < 0.05 were considered statistically significant.

### Classification Analysis

To assess the predictive utility of the identified FC-Traits, we trained a series of standard classifiers to predict a participant’s group status (Psilocybin vs. Placebo) (see Figure 1C). The subject-specific weights from any FC-Trait that showed a significant association with an external variable in the preceding regression analyses were used as predictive features. To ensure an unbiased performance evaluation, we employed a 10-fold stratified cross-validation framework. Within this framework, we trained and evaluated a standard set of machine learning classifiers: logistic regression, a linear support vector machine (SVM), and k-nearest neighbours (k=5). Model performance was quantified using several standard metrics for classification evaluation, including mean accuracy, Area Under the Receiver Operating Characteristic Curve (AUC), precision, recall, and the F1-score (Fawcett, 2006), aggregated across all test folds.

