## Supplemental Information for "Unmixing the Psychedelic Connectome: Brain Network Traits of Psilocybin"

### Supplementary Information

#### Scheme of connICA

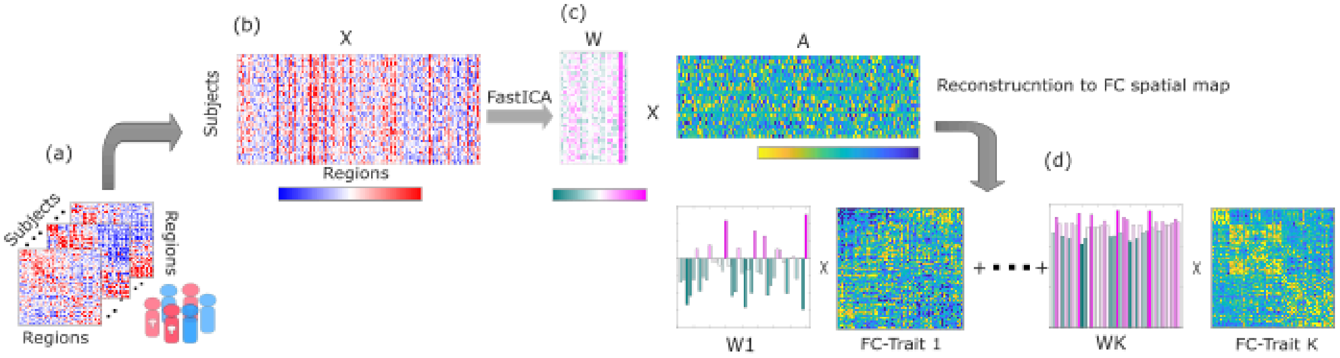


**Supplementary Figure 1. The Connectome Independent Component Analysis (connICA) Workflow.** This figure illustrates the core data processing and decomposition pipeline of the connICA framework. **(a)** Subject-specific 200*200 functional connectomes (FCs) are generated for all participants. **(b)** For each participant, the unique connectivity values (upper-triangular entries) are vectorized and concatenated into a single data matrix, X, with dimensions [subjects * connections]. This is performed without any prior grouping of participants. **(c)** The FastICA algorithm decomposes X into a subject-weight matrix, W [subjects * components], and an independent-component matrix, A [components * connections]. **(d)** Each row of A is reshaped back into a 200*200 matrix, representing a shared, system-level pattern of connectivity termed an "FC-Trait." The corresponding column in W provides the subject-specific weights, quantifying how strongly each individual expresses that particular trait.

**Supplementary Note. Calculation of Explained Variance for Optimal Model Selection**

To objectively quantify the goodness-of-fit for each set of robust components (RCs) identified within a given subsampling iteration and for a specific model order (e.g., 30 ICs), we implemented a reconstruction and regression procedure. This method assesses how well the discovered components can explain the variance in the original functional connectivity data.

The procedure was as follows: First, for each of the K robust components identified for a given model, a reconstructed connectome was generated. This was achieved by calculating the outer product of the component's mean subject-weight vector (a vector of size 40 * 1) and its spatial vector (a vector of size 1 * 19,900). This operation resulted in K separate reconstructed data matrices, each with the dimensions [40 subjects * 19,900 connections].

Second, these K matrices were each vectorized and then concatenated to form a final predictor matrix with dimensions [176,000 observations * K predictors], where each column represents the full reconstructed data from a single robust component.

Finally, a single multivariate linear regression model was fitted. The dependent variable in this model was the original vectorized data matrix for the 40 subjects in the subsample. The independent variables were the K columns of the predictor matrix. The Adjusted R-squared value from this regression model was then extracted. This single value represents the proportion of variance in the original connectivity data that is explained by the set of robust components for that specific model order, adjusted for the number of predictors.

**Supplementary Table 1. Stability and Composition of Robust FC-Trait Clusters**

| No. | ClusterID | Robustness Score |
|---|---|---|
| 1 | 73 | 99 |
| 2 | 64 | 97 |
| 3 | 2 | 94 |
| 4 | 59 | 94 |
| 5 | 74 | 94 |
| 6 | 92 | 92 |
| 7 | 93 | 89 |
| 8 | 77 | 88 |
| 9 | 89 | 88 |
| 10 | 101 | 88 |
| 11 | 80 | 86 |
| 12 | 86 | 85 |
| 13 | 110 | 84 |
| 14 | 88 | 81 |
| 15 | 81 | 76 |
| 16 | 98 | 72 |
| 17 | 39 | 68 |
| 18 | 78 | 65 |
| 19 | 71 | 64 |
| 20 | 72 | 64 |
| 21 | 87 | 62 |

**Supplementary Table 1. Details of the 21 Robust FC-Trait Clusters Identified via Hierarchical Clustering**. The table lists the final 21 clusters that surpassed the 60% robustness threshold. The "Robustness Score" quantifies the stability of each trait across the participant subsamples; a score of 99, for example, indicates that the spatial pattern represented by that cluster was identified as an optimal component in 99 of the 100 independent resampling iterations. The two traits found to be functionally significant in the main text correspond to specific clusters listed here: the Visual Divergent Thinking-Associated Functional Trait (VDT-FT) is the centroid vector of Cluster ID 64, and the Psilocin-Associated Functional Trait (PA-FT) is the centroid vector of Cluster ID 71.

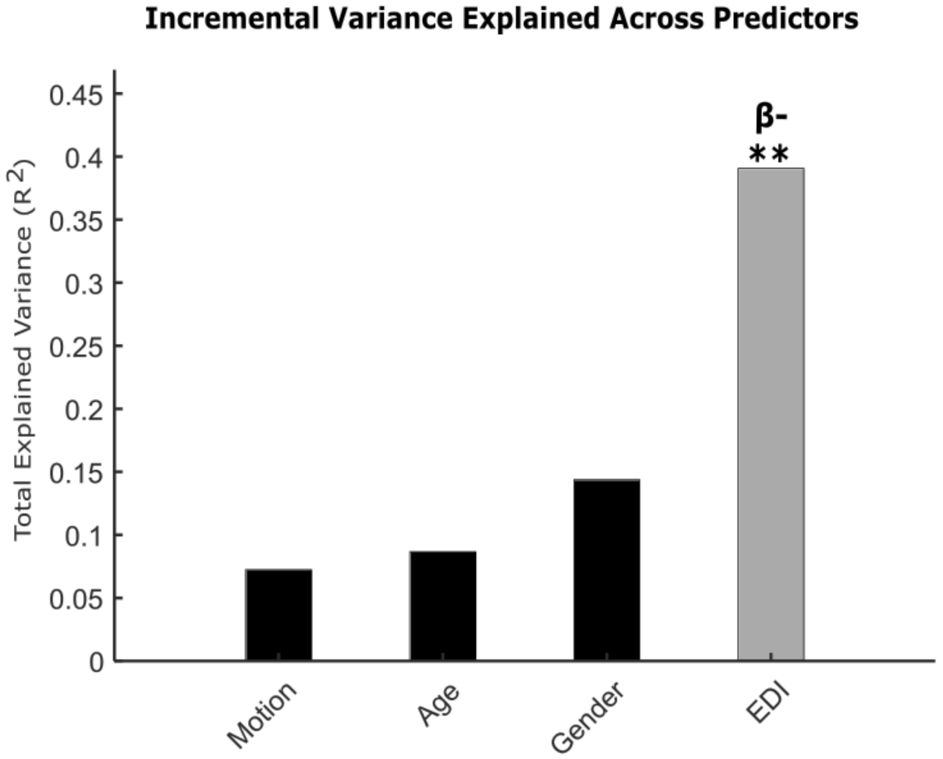


**Supplementary Figure 2. Association between PA-FT expression and Ego Dissolution.** The bar chart illustrates the incremental R-squared from the multiple linear regression model predicting PA-FT weights. After accounting for covariates (Motion, Age, Gender), the Ego Dissolution Inventory (EDI) score emerges as a significant predictor, with the full model explaining approximately 39% of the variance ($R^2 \approx 0.39$). The asterisks (**) denote the statistical significance of the EDI score's contribution to the model (uncorrected $p < 0.005$), while the negative beta sign (β-) indicates that higher subjective ratings of ego dissolution are associated with weaker expression of this trait.

**Note on the PA-FT, Psilocin, and EDI Associations:** We defined the PA-FT by its primary link to the objective, psilocin concentration rather than the subjective EDI score, as the EDI score showed collinearity with psilocin ($r \approx 0.6$) and reflects a downstream phenomenological effect. The strong secondary correlation with EDI is not a contradiction but a functional validation, as the trait's DMN-related spatial patterns are the well-established neural correlate of ego dissolution.

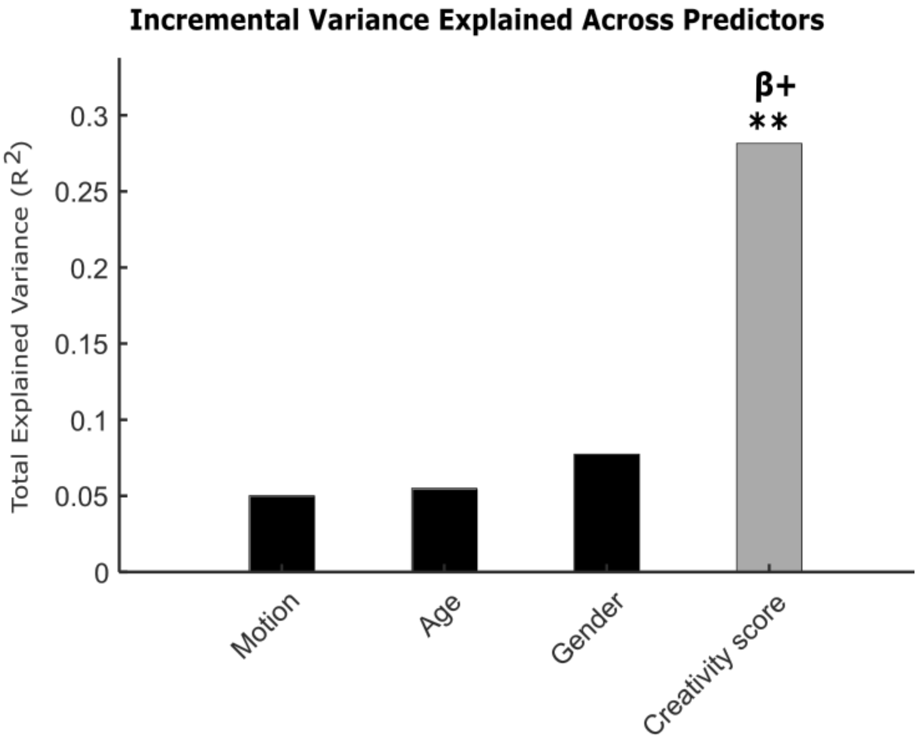


**Supplementary Figure 3. Association between VDT-FT expression and PCT originality score.** The bar chart illustrates the incremental R-squared from the multiple linear regression model predicting VDT-FT weights. After accounting for covariates, the Picture Concept Task (PCT) originality score emerges as a highly significant predictor, with the full model explaining approximately 28% of the variance ($R^2 \approx 0.28$). The asterisks (**) denote the statistical significance of the creativity score's contribution to the model (uncorrected $p < 0.005$), while the positive beta sign (β+) indicates that higher creative originality is associated with stronger expression of this trait.
